# Clinical and Translational Neuroscience

**DOI:** 10.1101/543678

**Authors:** Tom Gilbertson, Mark Humphries, J. Douglas Steele

## Abstract

In monogenetic generalized forms of dystonia, *in vitro* neurophysiological recordings have demonstrated direct evidence for abnormal plasticity at the level of the cortico-striatal synapse. It is unclear whether similar abnormalities contribute to the pathophysiology of cervical dystonia, the most common type of focal dystonia. We investigated whether abnormal cortico-striatal synaptic plasticity contributes to abnormal reward-learning behavior in patients with focal dystonia. Forty patients and forty controls performed a reward-gain and loss-avoidance reversal learning task. Participant’s behavior was fitted to a computational model of the basal ganglia incorporating detailed cortico-striatal synaptic learning rules. Model comparisons were performed to assess the ability of four hypothesised receptor specific abnormalities of cortico-striatal long term potentiation (LTP) and Long Term Depression (LTD): increased or decreased D1:LTP/LTD and increased or decreased D2: LTP/LTD to explain abnormal behavior in patients. Patients were selectively impaired in the post-reversal phase of the reward task. Individual learning rates in the reward reversal task correlated with the severity of the patient’s motor symptoms. A model of the striatum with decreased D2:LTP/ LTD best explained the patient’s behavior, suggesting excessive D2 cortico-striatal synaptic depotentiation could underpin biased reward learning in patients with cervical dystonia. Reversal learning impairment in cervical dystonia may be a behavioural correlate of D2 specific abnormalities in cortico-striatal synaptic plasticity. Reinforcement learning tasks with computational modeling could allow the identification of molecular targets for novel treatments based on their ability to restore normal reward-learning behavior in these patients.

## 1.0 Introduction

Cervical dystonia is a form of primary focal dystonia characterized by involuntary muscle contractions and posturing of the neck region that leads to significant disability and pain. The degraded action selection which underpins these involuntary movements have long been thought to be the consequence of dysfunction within the basal ganglia, based on clinical observation (Bhatia & Marsden, 1994; Berardelli *et al*., 1998), neuroimaging (Naumann *et al*. (1998); (Colosimo *et al*., 2005; Draganski *et al*., 2009; Zoons *et al*., 2011) and its secondary association with “extra-pyramidal” disorders (Burke *et al*., 1982; Rivest *et al*., 1990; Louis *et al*., 1999). The recent identification of families with novel mutations in genes which encode proteins richly expressed within the striatum implicated in dopaminergic signal transduction e.g. GNAL (Kumar *et al*., 2014), and ANO3 (Charlesworth *et al*., 2012) has reignited interest in “old ideas” about dopaminergic dysfunction being a common pathophysiological hallmark (Goodchild *et al*., 2013). For patients with cervical dystonia, detailed molecular and neurophysiological understanding of the nature of this derangement in signaling is crucial for the development of new treatments.

By its opposing effects on cortico-striatal plasticity at D1 and D2 receptors (Surmeier *et al*., 2007) dopamine determines which actions are selected for a specific context, by encoding action values in cortico-striatal synaptic strengths (Samejima *et al*., 2005). If dopamine signaling is abnormal in patients with cervical dystonia, one potential cause for the breakdown in action selection is the downstream effect of abnormal signaling on synaptic connectivity at the cortico-striatal synaptic interface (Peterson *et al*., 2010; Quartarone & Pisani, 2011). Abnormalities in striatal plasticity are thought to be a pathogenic mechanism common to all forms of dystonia (Peterson *et al*., 2010). In animal models of rare, childhood onset generalized dystonia (DYT1, DYT6), highly specific cortico-striatal plasticity abnormalities have been observed *in vitro* (Pisani *et al.,* 2006; Martella *et al*., 2009; Maltese *et al*., 2017; Zakirova *et al*., 2018). Whether defective cortico-striatal plasticity contributes to the expression of the common late onset focal forms, including cervical dystonia, remains unknown.

We therefore developed an experimental paradigm and computational model that aimed to test two hypotheses: i) cervical dystonia is associated with a measurable bias in reward-learning; ii) a specific abnormality in cortico-striatal plasticity could explain this abnormal behavior.

## 2.0 Materials and methods

### 2.1 Participants

The study was approved by the local Ethics Committee (East of Scotland Research Ethics Service, reference number 2014NG03) and written informed consent obtained from all volunteers. All patients were recruited from movement disorder clinics from four regional neuroscience centers in Scotland (Aberdeen, Dundee, Edinburgh and Glasgow). The diagnosis of cervical dystonia was made by a Consultant Neurologist with a specialist interest in movement disorders. Patients were all receiving botulinum toxin injections and behavioral testing was performed to coincide with maximal treatment response (2-8 weeks). The principal exclusion criteria for the study were: secondary dystonia and drug induced dystonia and previous or ongoing mood, anxiety or other psychiatric disorders. Patients taking anti-cholinergic or other centrally acting pharmacological agents for treatment of their cervical dystonia were excluded. Age, IQ and sex-matched controls were university and NHS staff members with no previous history or active symptoms of neurological or psychiatric disease.

#### 2.1.1 Rating scales

Clinical rating of cervical dystonia severity was assessed using the Cervical Dystonia Impact Profile [CDIP-58] (Cano *et al*., 2004). Ratings of mood, anxiety and obsessive-compulsive symptoms were derived using the HADS and Y-BOCS assessments (Zigmond & Snaith, 1983; Goodman *et al*., 1989). The NART was used to estimate IQ (Nelson HE, 1991).

#### 2.1.2 Behavioral task

Patients and controls performed a probabilistic reward (Figure 1) and loss (Supporting information Figure 1) reversal learning task based on previous reinforcement learning studies (Pessiglione *et al*., 2006; Gradin *et al*., 2011). Subjects were asked to learn from trial and error in order to obtain “vouchers” and were informed at the beginning of the task that their voucher score would be converted into money on completion. This ranged from £20-30 depending on the number of vouchers obtained. During the task one set of two pairs of fractal images were presented on a computer screen. Subjects chose one of the two fractals with a button press with the aim of maximizing wins (positive reinforcement) and minimizing losses (negative reinforcement) by trial and error. Trials where responses were not obtained within 2.5 seconds were not included in the analyses. Feedback following image choice informed subjects as to whether they “won” or “lost” a voucher (in the reward and loss trials respectively) or “nothing” where no change in score occurred. Reward and loss trials were presented randomly with an 80:20 probability of a reward or loss being the outcome with a total of 120 trials per task being presented. After 60 trials the contingencies were reversed, requiring subjects to extinguish their previously learned action-outcome relationship. The task was performed over three sessions with short between-session breaks and the reversal occurring midway through the second session. A five minute training session was performed before formal testing began.

**Figure 1:**
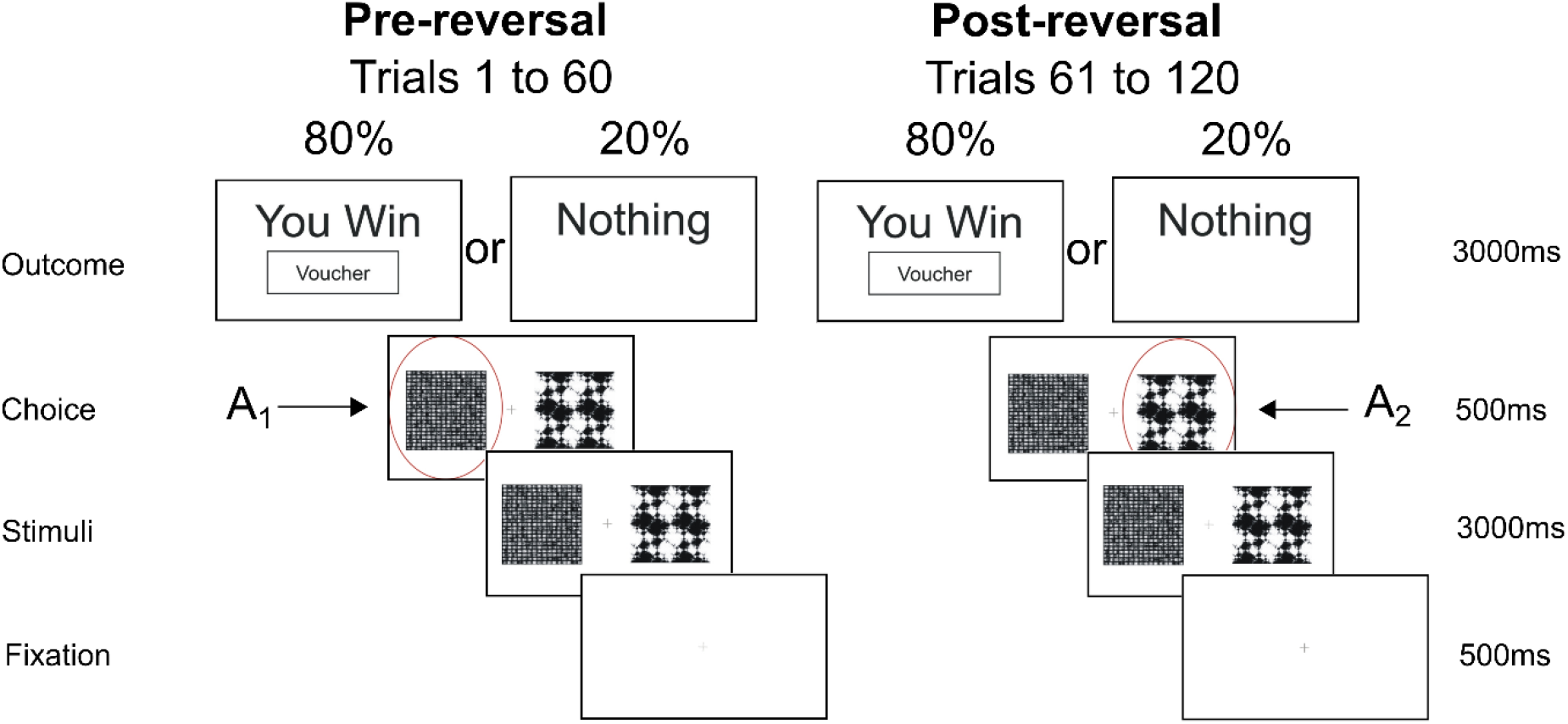
Probabilistic reversal learning task. Example of fractal images presented during a single reward trial. The probability of receiving a reward “voucher” reverses after 60 trials requiring the participant to suppress the previously learnt choice and learn to “reverse” their decision to choose the previously low value (pre-reversal) fractal. In the pre-reversal phase choice of A_1_ is associated with an 80% outcome of reward. Post-reversal this reduces to 20% with the choice of A_2_ being associated with 80%

The motivation behind our choice of task was the recognized role of the basal ganglia and in particular the striatum in reversal learning (Cools *et al*., 2002; Bellebaum *et al*., 2008). Reversal learning involves the initial acquisition and subsequent (post-reversal) extinction of action-outcome contingencies. These processes depend upon both direct (D1) and indirect (D2) pathway medium spiny neurons (MSN’s) (Cools *et al*., 2009; Nakanishi *et al*., 2014; Cox *et al*., 2015). By using a task which engages both striatal populations, our aim was to elucidate specific isolated or combinations of abnormalities, in both D1 and D2 plasticity. The paradigm was programmed using MATLAB (MathWorks, Natick, MA) with PsychToolbox (Brainard, 1997).

### 2.2 Functional MRI data acquisition and image analysis

Functional MRI was acquired in twenty of the controls and nineteen of the patients performing the task (one patient was unable to tolerate scanning due to claustrophobia, but completed the behavioural experiment outside the scanner). For each participant, functional whole-brain images were acquired using a 3T Siemans Magnetom TimTrio Syngo scanner using an echoplanar imaging sequence with the following parameters: angle = 90°, field of view = 224 mm, matrix = 64 × 64, 37 slices, voxel size 3.5 × 3.5 × 3.5 mm. Images where visually inspected for artefacts and preprocessing was performed in SPM8 (http://www.fil.ion.ucl.ac.uk/spm). Images were realigned and co-registered to the SPM8 Montreal Neurological Institute echo planar imaging template. The average, realigned co-registered image for each subject was used as a template to normalize each realigned and co-registered volume to the SPM8 echo planar imaging template image before smoothing. For first level analysis, an event related model-based analysis was implemented with onsets at the outcome time (when the subject received feedback as to whether their choice was sucessful or not). The reward prediction error signal generated from the individual subjects optimally fitted ‘Standard Reinforcement Learning Mode)’ (see below) was used to parametrically modulate a truncated delta function convolved with the haemodynamic response function. Second level analysis was restricted to one-group t-tests to assess for significant activation patterns within groups and two-group t-tests for estimates of between group (patient versus controls) differences in RPE activation patterns. Using a popular Monte-Carlo method (Slotnick et. al., 2003), significance was defined as P<0.01 at a whole brain, Family-Wise Error corrected level, achieved by a simultaneous requirement for a p<0.05 voxel threshold and a cluster extent >120 voxels.

### 2.3 Computational modelling

Two models were fitted to the observed behaviour. In the first instance a standard reinforcement learning model was chosen (Sutton RS, 1998) which robustly captures phasic dopamine neuron firing in the form of a reward prediction error signal (RPE). The dynamics of the RPE signal in this model is determined by estimating two parameters, the learning rate (*α*) and reward sensitivity (*β*). This type of model has been used previously to explore abnormal presynaptic phasic dopamine signaling in pathological states such as Parkinson’s disease (Peterson *et al*., 2009). In contrast, presynaptic signaling is thought to be preserved in cervical dystonia (Naumann *et al*., 1998), so our aim was to test the hypothesis that post-synaptic cortico-striatal plasticity was abnormal. To that end we developed a detailed basal ganglia model (Figure 2A) which combined phasic dopamine signaling with cortico-striatal synaptic plasticity dynamics (Gurney *et al*., 2015). This model required estimation of six parameters: learning rate (*α*) and reward sensitivity (*β*), plus four “plasticity coefficients” (*a*_1_,*b*_1_,*a*_2_,*b*_2_). Each coefficient determined the relative influence of phasic dopamine release on the cortico-striatal synaptic weight; with *a*_1_,*b*_1_ scaling D1-LTD and D1-LTP, and *a*_2_,*b*_2_ scaling the D2 LTP and D2-LTD synaptic changes respectively (Figure 2A).

**Figure 2:**
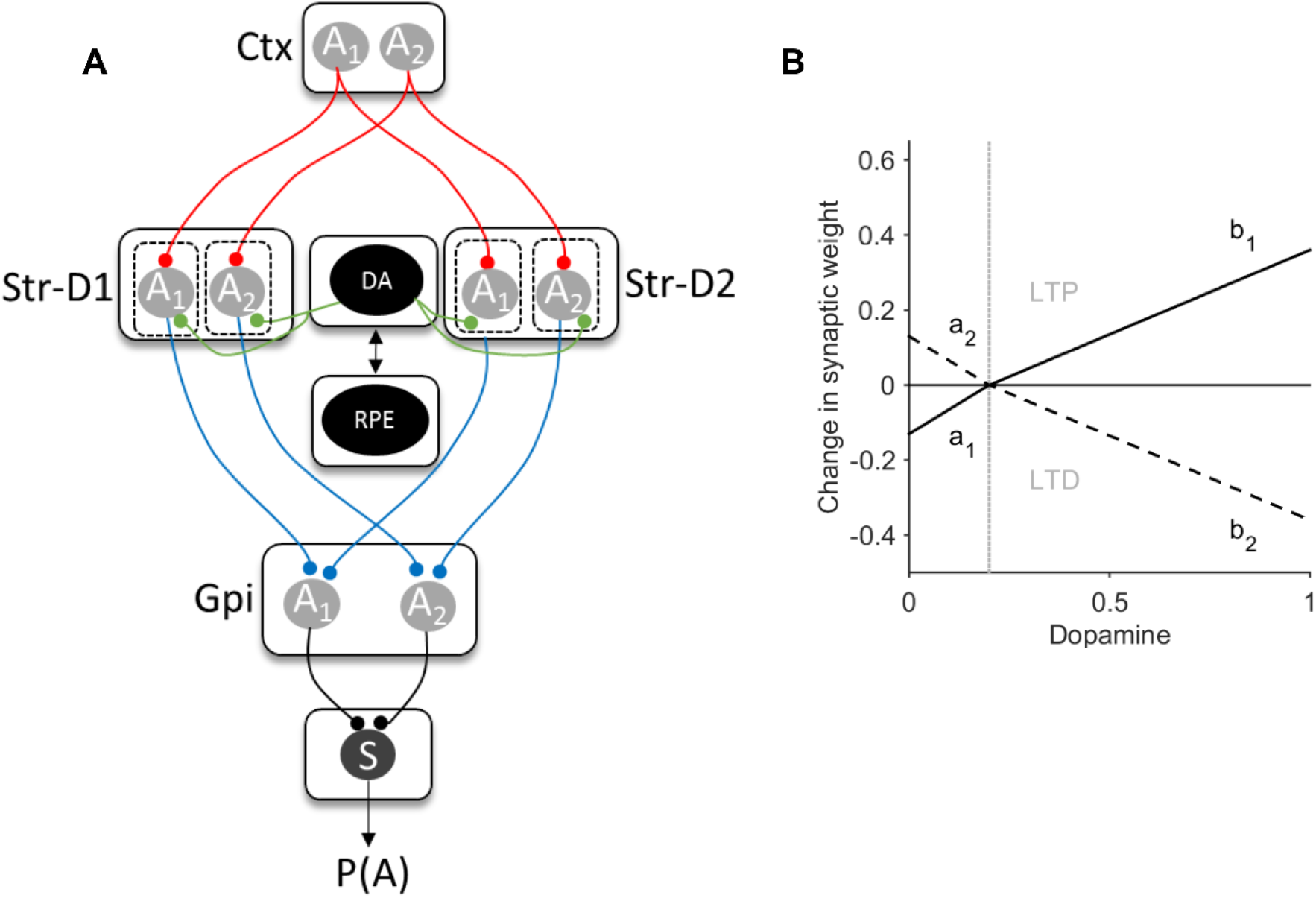
Basal ganglia plasticity model. **(A)**. Schematic model architecture. Nuclei include the direct pathway striatal neurons (Str-D1), indirect pathway (Str-D2) and globus pallidus interna (GPi). Grey circles represent the available two actions (A1 and A2) represented in the cortex (Ctx) and subcortical nuclei. Red lines represent excitatory connections, blue inhibitory. Green represents the neuromodulatory influence of dopamine on the striatum. Functions critical to the model include the dopamine-weight change function (DA), the reward prediction error signal (RPE) and the softmax (S) equation. The value of an action is assumed to be represented in the cortex (Ctx) for one of two actions (A1 and A2). The striatal output is determined by the product of the cortico-striatal synaptic weight and the cortical input for the action. The NoGo activity of one action was subtracted from its “Go” D1” self (ie. Gpi(A1)=StrD1(A1)-StrD2(A1). The pallidal activity for each action is then converted into a choice probability P(A) by entering this into the softmax function (S). **(B)** The expected change in synaptic weight as a function of dopamine released is represented by the dopamine-weight change curve (DA). The four plasticity coefficients (a_1_,b_1_, a_2_,b_2_) in the model determine the gradients of this function, which in turn govern the magnitude of LTP and LTD at the cortico-striatal synapse. The solid line represents the D1 dopamine weight changes, the dashed line the D2 equivalent. The grey vertical line represents the point of inflection at which dopamine is considered above or below baseline levels, corresponding to phasic “bursts” or “dips” in the reward prediction error signal

#### 2.3.1 Standard Reinforcement Learning model

The SARSA model (Sutton & Barto, 1998) seems more consistent with actual dopamine firing than alternative temporal difference models (Niv et al 2006) and robustly captures the dynamics of phasic dopamine neuron firing in the form of a reward prediction error signal (RPE) represented by (*R*(*t*) − *Q*(*A, t* − 1)) in the equation;

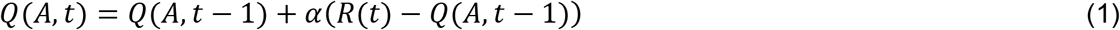

where the Q-value of action A in on trial t is the Q-value of the action on the previous trial *(t−1)* updated by the RPE. R_t_ is the outcome (reward[1] or nothing[0]) and *α* is the learning rate. The choice of action in the trial-by-trial reinforcement learning model is determined by the sigmoid or softmax function, where the probability *P* of choosing action A_1_ over action A_2_ is defined as:

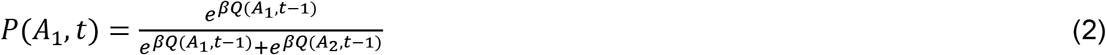

with *β* the reward sensitivity or inverse temperature parameter. Parameters where estimated using a random effects, expectation maximization (EM) procedure implemented in Matlab code provided by QJM Huys (https://www.quentinhuys.com/tcpw/code/emfit/).

To test for differences in the modelled RPE signal, individual subject estimates of the learning rate (*α*) and reward sensitivity (*β*) for patients and controls were generated. These were then used to test the null hypothesis of no significant difference in the model parameter estimates between groups. Comparative statistics were performed on normalized parameter estimates (Huys *et al*., 2013) where the normalized learning rate is a normally distributed random variable with mean −1 and related to the learning rate, *α*, by the following:

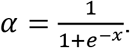

#### 2.3.2 Basal Ganglia model

A model was developed which incorporated a suitable level of detail to allow inference of striatal synaptic plasticity abnormalities, whilst being simple enough that it could be fitted to behavioral data. This “basal ganglia” model was designed to extend any general conclusions, inferred from a trial-by-trial reinforcement learning model, into more detailed mechanistic abnormalities that could be related to dopamine’s role in biasing action selection in the basal ganglia.

At the core of the basal ganglia model (Figure 2A) was a hybrid approach, incorporating standard reinforcement learning rules (to produce the RPE from equations 1 and 2) with cortico-striatal synaptic plasticity parameters, derived from the model of Gurney *et al*., 2015. Aiming for a simple parsimonious model, two striatal “populations” were assumed *S_D1_* and *S_D2_*; the D1 receptor expressing direct and D2 receptor expressing indirect pathways respectively. The striatal activity of each population *n*, on trial *t* for action *A* was:

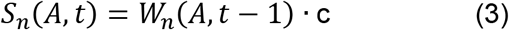

where *W* is the cortico-striatal synaptic weight and c is a constant input of 1. Here we use a model with two actions (*A*_1_, *A*_2_) corresponding to the choice between the two options.

Competition between the two striatal pathways for control of basal ganglia output was of most interest. Thus, the pallidal output for action *A*_1_ was defined as:

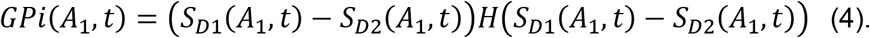

where *H()* is the Heaviside step function: *H(x)* = 0 if *x* ≤ 0, and *H(x)* = 1 otherwise; and similarly for action *A*_2_. The pallidal output was then proportional to the amount of disinhibition of the basal ganglia’s targets. In turn the probability of choosing action A_1_ was determined by the softmax equation with the basal ganglia’s output substituted for the value term:

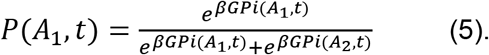

The cortico-striatal synaptic weights are assumed to take on only positive values and are modified at the synapse corresponding to the chosen action *A*

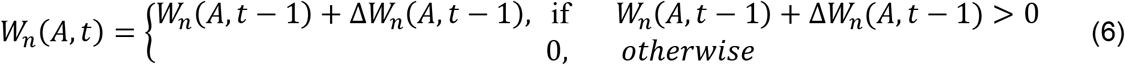

The change in synaptic weight is assumed to obey a Hebbian two-factor rule as the product of the striatal postsynaptic activity (S) and the neuromodulatory influence of dopamine (Δ*d_n_*(*t*)) assuming constant (pre-synaptic) cortical input:

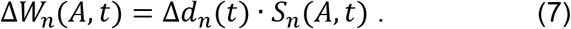

The extent of dopamine related changes in synaptic strength are governed by separate D1 and D2 functions following the detailed model of dopamine-modulated plasticity in Gurney *et al*., [their Fig. 13], and capture the dependence of the sign and magnitude of synaptic change on both the concentration of dopamine (DA) and the receptor-type of the post-synaptic striatal neuron (D1 or D2).

For the general case of a *D_n_* dopamine receptor subtype, the magnitude Δ*d_n_* of dopamine’s effect on synaptic plasticity is,

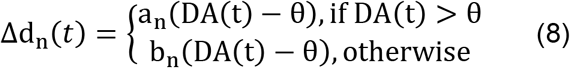

where (*a_n_,b_n_*) are coefficients determining the dependence of synaptic plasticity on the current trial’s level of dopamine DA(t), and the constant *θ* determines the baseline level of dopamine. Following the predictions of Gurney *et al*., 2015, for D1 neurons we expect here (*a*_1_,*b*_1_) > 0, so that dopamine levels below baseline cause a decrease in cortico-striatal weight (long term depression; LTD), and dopamine levels above baseline cause an increase in weight (long term potentiation; LTP); for D2 neurons we expect here (*a*_2_,*b*_2_) < 0, thus giving LTP below baseline, and LTD above it. However, in fitting this model to the behavioral data, we did not restrict the sign of the coefficients.

Following standard RPE accounts of dopamine signaling^9^, values of DA>*θ* are phasic increases, a positive prediction error, whereas DA<*θ* represent phasic dips in dopamine, the negative prediction error signal. Thus the RPE(t) signal of Equation 1 was mapped to DA(t) in Equation (9) as:

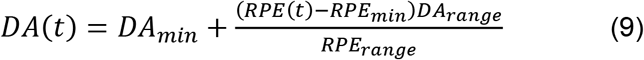

To map the RPE(t) signal in the range [−1,1] to the DA value in [0,1] around the baseline of *θ*, we use: if RPE(t) < 0, *DA_min_* = 0, *DA_range_* = *θ*, *RPE_min_* = −1, *RPE_range_* = 1; otherwise *DA_min_* = *θ, DA_range_* = 1 − *θ, RPE_min_* = 0, *RPE_range_* = 1.

The baseline level of dopamine was set at a value of *θ* = 0.2 for all simulations. Initial values of the synaptic weights were set to 0.2 and action values value at 0.5, leaving the four plasticity parameters (*a*_1_,*b*_1_,*a*_2_,*b*_2_), and *α* and *β*, from the equations (1) and (2), to estimate.

The four plasticity parameters govern the magnitude of D1-LTD (*a*_1_), D1-LTP (*b*_1_), D2-LTP (*a*_2_) and D2-LTD (*b*_2_). Figure (2B) gives an illustrative example of the interaction between the dopamine levels (as governed by the RPE) and its effect on striatal synaptic strength using the default plasticity coefficient parameters from Gurney *et al*. 2015,.

#### 2.3.3 Basal ganglia model fitting – RPE parameters

At all stages of the fitting procedure, parameter optimization was performed by minimizing the negative log-likelihood of the data given different values of the model’s parameters. Our initial approach was to define a realistic “physiological” parameter space by fitting the control subjects’ behaviour. As we wanted to eliminate any influence of the Q-learning / RPE parameters on the eventual patient fits, we first estimated individual learning rate (*α*) and reward sensitivity (*β*) parameters for the control subjects whilst keeping the four plasticity parameters, *a*_1_,*b*_1_,*a*_2_,*b*_2_ fixed at default values 0.5,0.3,−0.5,−0.5 respectively. For each control subject, *α* and *β* estimates were derived following a grid search over the explored ranges of [0, 0.5] and [0, 0.1] respectively. The parameters derived from this grid search were then used as starting values for further optimization using Matlab’s *fmincon* function. The purpose of this initial fitting procedure to the control subjects was to find values of *α* and *β* which could then be used as constants for all subsequent estimation of the plasticity coefficients. All subsequent fitting of the plasticity parameters for each subject was therefore performed whilst using this average value of *α* and *β* as constants. This approach was used to prevent overfitting of the behavioural data and ensure that the model fitting resulted from optimization of a four parameter search for striatal plasticity parameters, to test our hypothesis, rather than parameters relating to phasic dopamine release.

#### 2.3.4 Basal ganglia model fitting – plasticity parameters

An exhaustive search over the entire parameter space was done as defined by the four plasticity parameters. The aim was to eliminate biologically unfeasible or redundant parameter space to define meaningful bounds of the constrained fitting that would follow. This approach was also found to significantly improve the consistency of parameter estimates by avoiding spurious local minima. As the “*a*”parameters are modified over a narrower range of dopamine, we allowed the search limits to be broader than the“*b*” parameters leading to ranges of [−2,2] and [−1,1] respectively. Given the four plasticity parameters could take on either positive or negative values within the parameter space, we divided this into 16 “quadrants,” each corresponding to a parameter combination with positive and negative signs. For example, the first quadrant with parameter signs [+, +, +, +] is thus defined by [0,0,0,0] for the lower bounds, [2,1,2,1] for the upper bounds. For each quadrant a coarse grid search on the parameters bounds was done followed by function minimisation with *fmincon*, using the course grid search minima as starting values and the quadrant’s bounds as *fmincon’s* constraints. The parameter space quadrant which produced the most reliable fits (lowest mean negative log likelihood scores) was the chosen for all subsequent analysis. This “exhaustive search” was performed for the control subjects only.

#### 2.3.5 Basal ganglia model fitting – hypothesis testing

Our aim was to establish whether pathological combinations of biologically feasible striatal plasticity abnormalities could explain differences in the patients’ behavior in the reversal learning task. Using the control estimates as normative data, parameter space bounds were defined which could be considered “pathological” and which could be used to test specific hypotheses on their ability to model patient’s behavior. Specifically, we chose four separate hypothetical combinations of plasticity deficits: H_1_) increased D1-LTP : D1-LTD; H_2_) decreased D1-LTP : D1-LTP; H_3_) increased D2-LTP : D2-LTD; H_4_) decreased D2-LTP : D2-LTD. These combinations were chosen as previous *in vitro* experiments have demonstrated a wide range of potential cortico-striatal plasticity abnormalities, but to date the receptor specificity of these remains unknown. With these four hypothetical combinations we could establish the most likely imbalance of cortico-striatal plasticity in a receptor specific manner. In order to define the parameter range that correspond to “pathological” hypothetical combinations, we fitted a non-parameteric kernel to the control estimates for each of the four parameters. We then derived a probability density function from this fit to obtain the 5% and 95% confidence limits for each parameter. These were then used to determine the bounds used for the fitting of the basal ganglia model to the patients’ behaviour. For example, in order to fit H1 (increased D1-LTP, decreased D1-LTD), the initial grid search was performed on a parameter space that was bounded between the 95% limit and 2 (the maximal parameter value) for “*a*_1_”; between 0 (the minimum value) and the 5% limit for “*b*_1_”, whilst “*a*_2_ “ and “*b*_2_” were constrained between the 5% and 95% confidence limit of the control parameter space. For each patient, the same procedure of grid search within the bounds defined by the hypothesis in question was followed by further optimisation of the parameter estimate using *fmincon*. As the number of parameters remained fixed across each hypothesis tested (and between model structures) we compared each combination of abnormal plasticity combinations by finding the model with the minimum negative log likelihood. This fitting procedure was performed separately for the reward and loss behavioural data.

### 2.4 Statistics

Behavioural and participant demographic data were tested for normality of distribution using Kolmogorov-Smirnov test. Normally distributed data were analysed using Student’s t-test with the Mann-Whitney U test was used as a non-parametric alternative where necessary. When more than one variable of interest was tested we used either one-way or multivariate ANOVA. In the case of categorical data, Chi-squared test was used.

## 3.0 Results

### 3.1 Demographics, rating scales and behavior

The reinforcement learning task was completed by forty patients with cervical dystonia (CD age = 55.2+/− 10.0 [range 28-72]; 28 female) and forty sex and age-matched controls (CTRL age 54.4 +/− 9.18 [range 26-74]; 27 female). The average cervical dystonia rating scale, (CDIP-58) score for the patients was 41 +/− 15 (range 22-80). Patients and controls were well matched with no significant difference in average IQ as indexed by the NART (CTL 118.18, +/− 3.8 CD 118.7 +/− 4.1,) or Y-BOCS scores (CTL 4.2 +/− 3.1, CD 5.1 +/− 4.3). There were numerically higher levels of anxiety and depression in the patients as indexed by the HADS-A (CTL 3.5+/− 3.1, CD 5.2+/2.596 two-tailed t-test, t(78) = 2.4, p=0.009) and HADS-D (CTL 1.42+/− 2.3, CD 2.66+/− 2.59 twotailed t-test, t (78)= 2.02, p=0.015) but this was not clinically significant and patients did not satisfy criteria for a mood or anxiety disorder. Patients and controls completed the same number of trials of both the reward and loss avoidance task (control 238+/− 1, patients 238 +/− 2.5).

Patients had a selective post-reversal deficit in reward-learning (Figure 3A). A one-way ANOVA demonstrated a significant between-group difference in the performance of the reward task (F(1,78) = 4.62.9, p = 0.03). The number of rewards won pre-reversal was similar between patients and controls, (CD = 50+/− 4.0, CTL 48+/− 3.2), Mann-Whitney z(78) = −1.61, p=0.11); however, the number of rewards won post reversal were significantly less for patients (CD = 30.1+/− 5.1, CTL= 38 +/− 4.5, Mann-Whitney z(78)= −2.47, p=0.013) implying an impairment in reward-based reversal learning (Figure 4A). No difference in the pre-or post-reversal learning performance in the loss avoidance task between patients and controls was observed (one way ANOVA between group difference F(1,68) = 1.15, p = 0.28), suggesting that learning from positive (reward) rather than negative (aversive) feedback was selectively impaired in patients with cervical dystonia (Figure 5).

**Figure 3:**
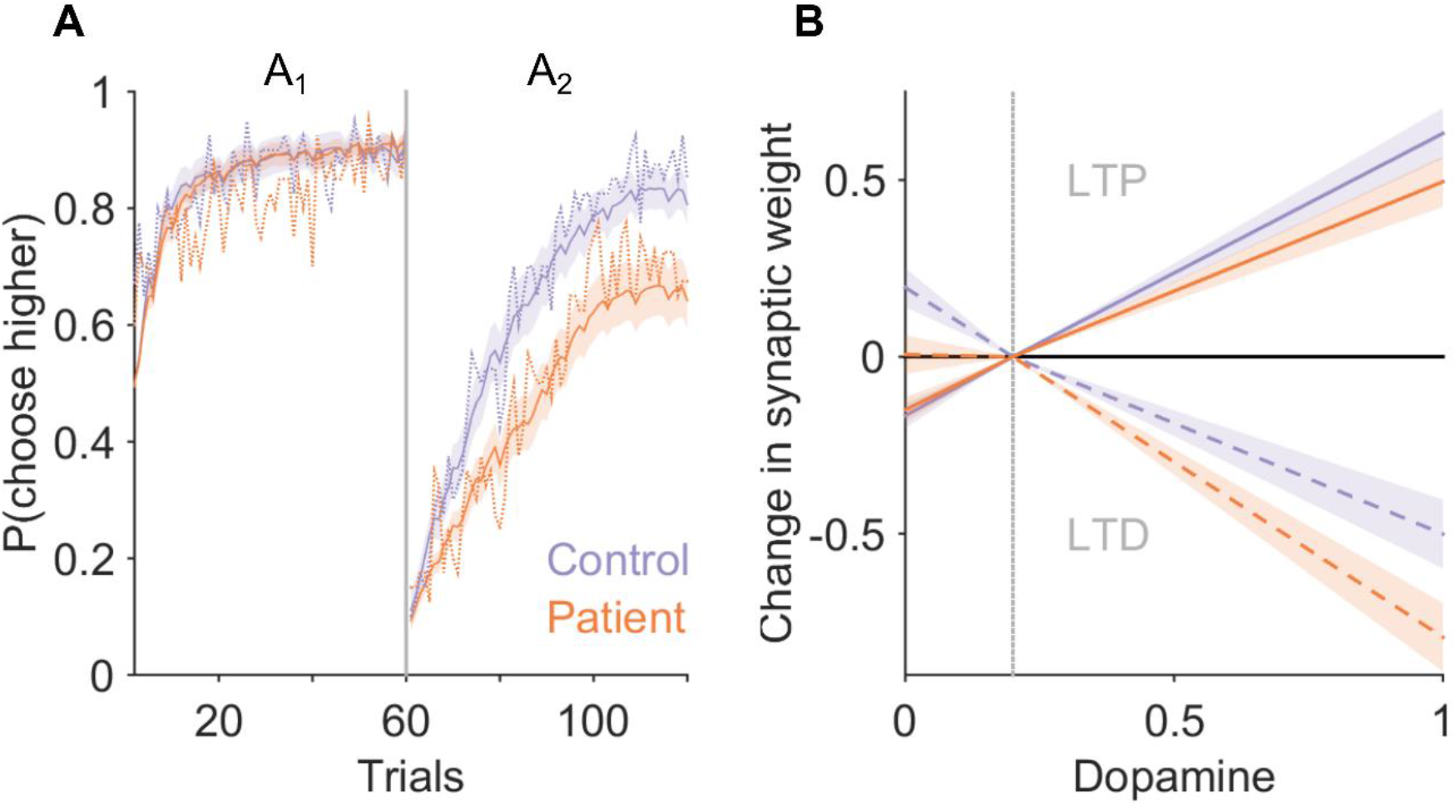
Impaired reward learning in cervical dystonia is explained by a model of the basal ganglia with abnormal D2 cortico-striatal plasticity. (A) Average probability of choosing high value actions “A1” and “A2” pre- and post-reversal reversal of contingencies in the reward task. Patient average choice probability in red, controls in blue. Vertical dotted line represents the point of contingency reversal. Dashed lines represent experimental behavior from patients and controls. The basal ganglia models performance following estimation of the optimal model parameters is superimposed (solid lines with 95% confidence limits represented by the shaded region). The models behavior closely overlaps with the experimental behavior of both the patients and controls. The number of rewards won post reversal in the reward task was significantly reduced in patients compared to controls (Mann-Whitney z(78)= −2.47, p=0.013). A basal ganglia model with increased D2-LTD : LTP explained the patients behavior best in 29 of the 40 patients fitted. (B) The average dopamine-weight change curve for the patients (red) and controls (blue) with 95% confidence limits represented by the shaded region. Solid lines, represent D1 and dashed lines, represent D2 curves respectively. During phasic “bursts” in dopamine the patients undergo significantly greater D2-LTD and correspondingly reduced D2-LTP during phasic “dips” below baseline (vertical grey dashed line).

**Figure 4:**
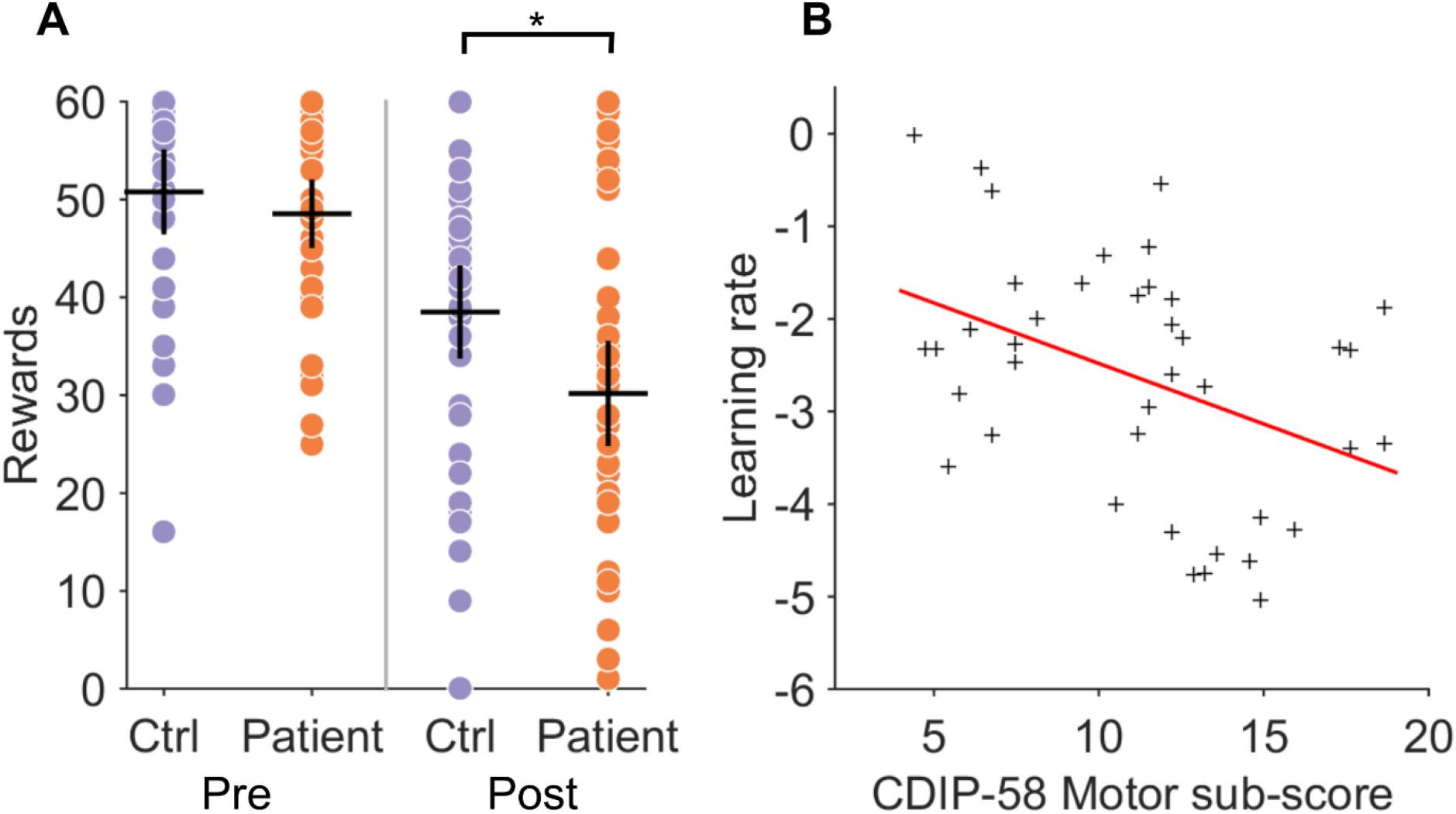
Cervical dystonia reward learning is impaired and correlates with severity of motor symptoms. (A) Individual performance in patients (red dots) and controls (blue) demonstrates significant reduction in number of rewards obtained post reversal (*p=0.013, Mann-Whitney z(78)= −2.47). Black crosses represent mean and 95% confidence limits. (B) The normalised learning rate (σ^(−1) α) estimate for each individual patient is plotted against their CDIP-58 motor sub-scores. There was a significant negative relationship between motor score severity and learning rate (R2 = 0.17, p = 0.008).

**Figure 5:**
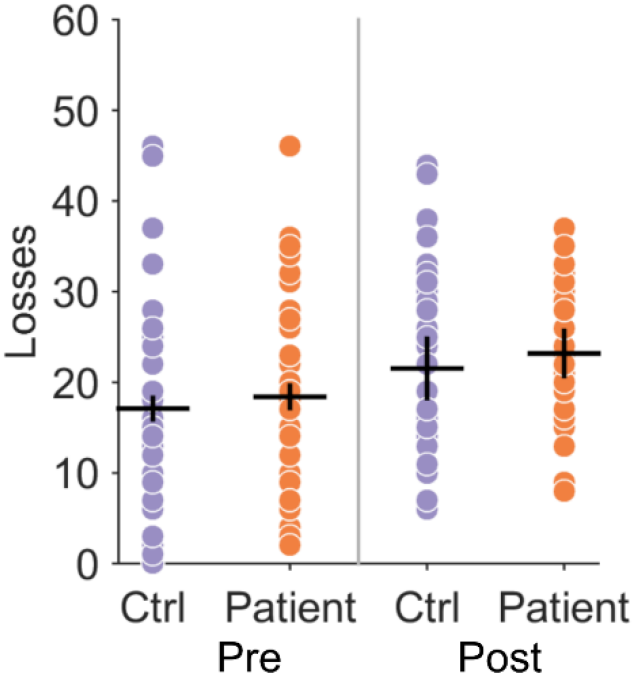
Loss avoidance learning in patients and controls. Individual performance in patients (red dots) and controls (blue) demonstrates no significance difference in loss avoidance performance. Black crosses represent mean and 95% confidence limits.

### 3.2 Reinforcement learning model fitting

To test the hypothesis that phasic dopamine signaling was abnormal in patients we modelled subject choice data from the reward behavioral task using a trial-by-trial reinforcement learning model^33^. This model was fitted to each individual participant’s behavioral choices in the reward task. A significant reduction in the learning rate parameter (α) was observed in the patients (CD α = −2.63+/− 1.29, CTL, α = −1.92 +/− 0.99, two-tailed t-test p=0.007, t(78)= −2.7) but not in the reward sensitivity parameter β (CD, β = 1.7 +/− 0.5, CTL, 1.8 +/− 0.56, t(78) = −0.7, p = 0.42). This is consistent with the patients placing less weight on the prediction error signal, as the learning rate parameter α multiplies the prediction error term in the trial-by-trial reinforcement learning model. No correlation was observed between individual values of α and the total CDIP-58 score (R^2^ = 0.07 p=0.13). However, a negative correlation between the “Head and Neck” symptom sub-score was present (R^2^ = 0.17, p = 0.008), meaning patients with the most severe motor symptoms tended to have the lowest learning rates (Figure 4B), and hence were most impaired at learning from positive feedback. We found no correlations with the learning rate parameter and the other CDIP-58 sub-scores, which include global estimates of disease impact on quality of life and psychosocial wellbeing, supporting a specific relationship between impaired reversal learning and motor symptoms in these patients. No correlation was found between the reward sensitivity parameter β and total CDIP-58 clinical rating scale or its sub-scores.

### 3.3 Event related fMRII analysis

Blunted reward prediction error signaling (as indexed by a lower learning rate) might be an explanation for the impairment in reward based learning in these patients and has been proposed for different patient groups. However, the results of an event-related model-based fMRI analysis of participants demonstrate that RPE signaling within the basal ganglia was intact in patients and at levels comparable to controls (Figure 6). Controls exhibited prediction error encoding within the striatum with maximal clusters in the left putamen, MNI co-ordinates [−10,10,−6], T-value = 5.14 and right caudate nucleus (MNI co-ordinates [10,7,−10], T-value = 5.08, both P <0.01, cluster extent corrected across the whole brain. There was no significant between group differences (at P = 0.05) suggesting comparable RPE encoding in patients and controls.

**Figure 6:**
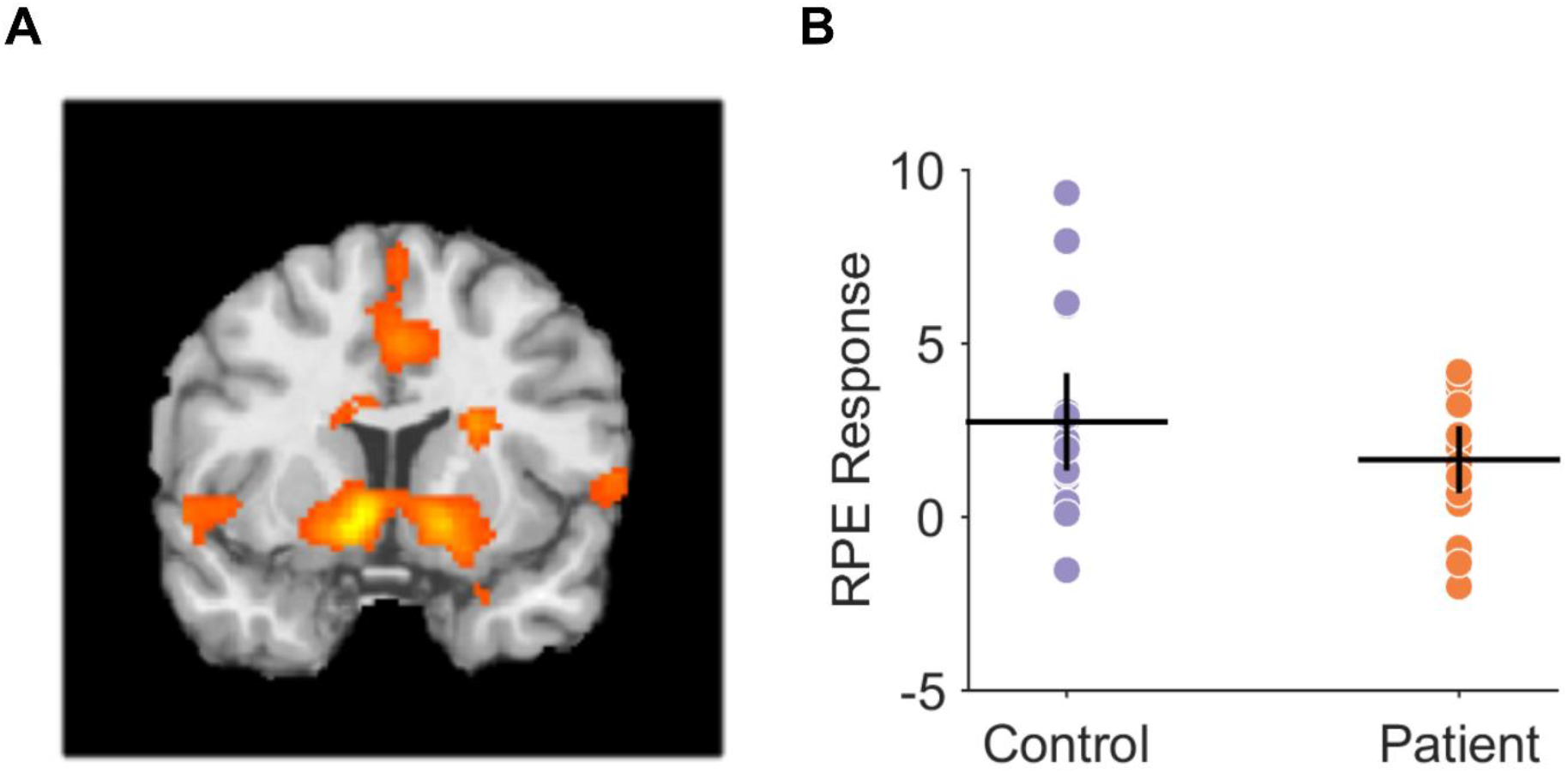
Reward prediction error encoding in controls and patients during reversal learning task. (A) Controls exhibited prediction error encoding within the striatum with maximal clusters in the left putamen, MNI co-ordinates [−10,10,−6], T-value = 5.14 and right caudate nucleus (MNI co-ordinates [10,7,−10], T-value = 5.08, both P <0.01, cluster extent corrected across the whole brain. There was no significant between group differences (at P = 0.05) suggesting comparable prediction error encoding in patients and controls. (B) Region of interest analysis from left putaminal RPE-fMRI signal in controls and patients with mean and 95% confidence limits supporting comparable levels of RPE signaling in both groups

### 3.4 Basal ganglia model fitting

#### 3.4.1 Controls

Our fMRI result suggests that (pre-synaptic) phasic dopamine signaling is normal in patients and a more likely explanation for impaired reversal learning is abnormal post-synaptic modification in cortico-striatal strength. To further understand this potential mechanism, the ‘Basal Ganglia’ model was used which, as above, incorporated standard reinforcement learning rules with details of cortico-striatal plasticity (Figure 2), allowing inference of synaptic weight changes pertaining to direct (D1) and indirect (D2) pathway potentiation (LTP) and depression (LTD). Six parameters were estimated; two relating to a standard reinforcement learning model (the learning rate (*α*) and reward sensitivity (*β*) parameters) and four plasticity coefficients representing D1-LTD, D1-LTP, D2-LTP and D2-LTD represented by (*a*_1_,*b*_1_,*a*_2_,*b*_2_) respectively (Figure 2B). All initial fitting was performed on the control behavior in order to define the “physiological” parameter space prior to testing hypothetical pathological explanations for the patients abnormal behavior. As the purpose of this model was to test for abnormal post-synaptic plasticity abnormalities, here we fixed the learning rate (*α*) and reward sensitivity (*β*) parameters, which govern the phasic dopamine signals dynamics, to constant values. The final estimated *α* and *β* values estimated for the 40 control subjects were *α* = 0.036 +/− 0.05, *β* = 0.28 +/− 0.14 for reward learning, and *α* = 0.11+/− 0.22, *β* = 0.36 =/− 0.24 for loss avoidance. All subsequent fitting of the model used these average values whilst allowing the remaining four plasticity parameters to vary. By excluding the patients from this analysis, this further reinforced the a *priori* assumption of our model fitting and results of the fMRI analysis, that phasic RPE signaling was intact and physiologically comparable to control levels, ensuring that any difference in behavior was explained by differences in plasticity parameters.

Next an exhaustive search over the entire parameter space was done defined by systematic variation of the four plasticity parameters eliminating biologically unfeasible or redundant parameter space in order to define the bounds of the constrained fitting that would follow. This analysis (Figure 7) demonstrated that only a small number of parameter combinations modelled the behavior well, the majority fitting the behavior poorly. Of the few parameter combinations which did model the behavior consistently well, only one was consistent with the known physiological plasticity modifications of dopamine at D1 and D2 synapses. We therefore settled on constraining the parameter bounds to within this territory of the parameter space where *a*_1_ = [0,2],*b*_1_ = [0,1], *a*_2_ = [−2,0],*b*_2_ = [−1,0].

**Figure 7:**
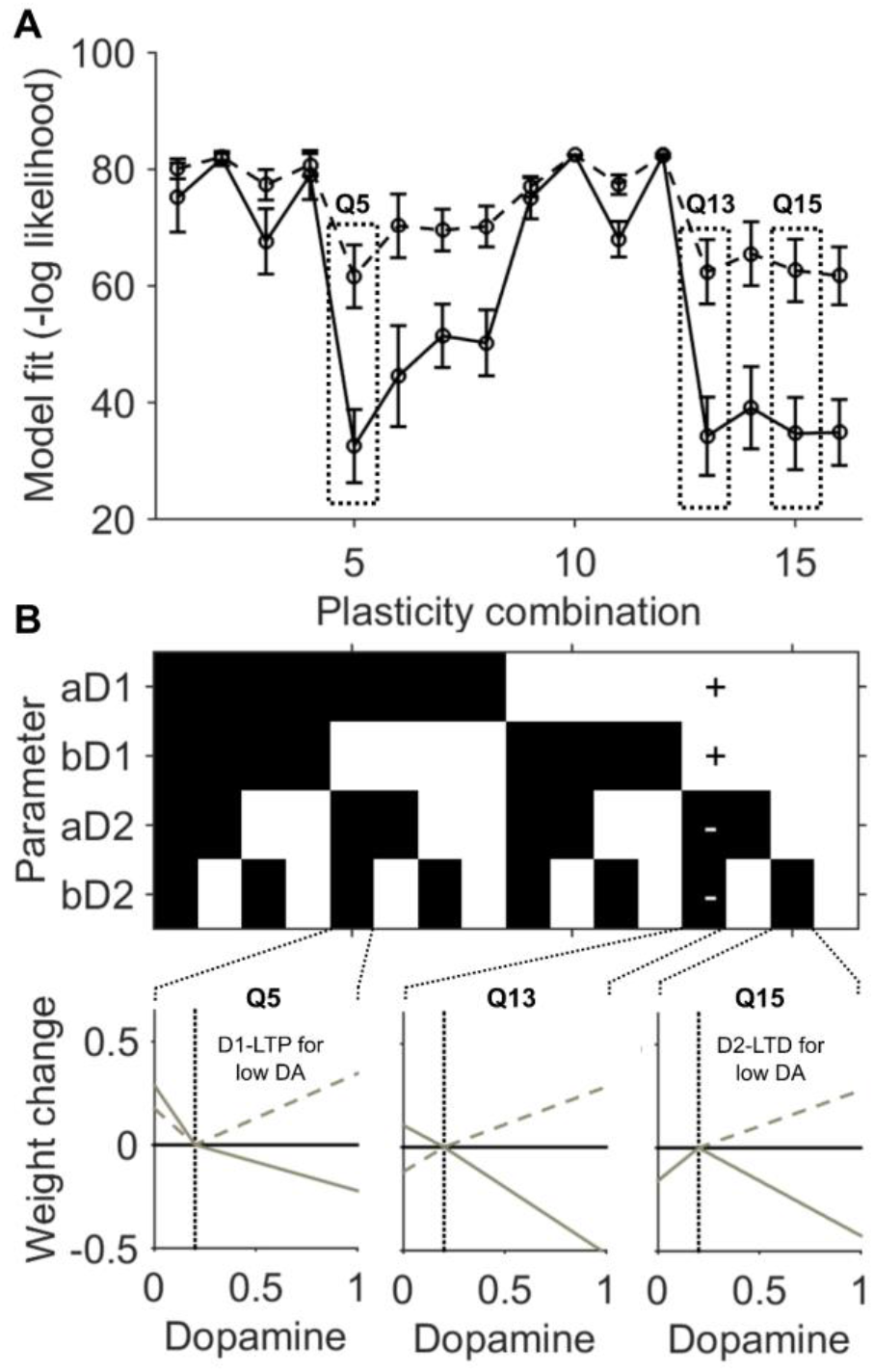
Quadrant parameter space mapping. (A) The average negative log likelihood (error bars represent 95% confidence intervals) for model fitting to the control behavior. Solid lines represents reward fitting, dashed lines results from fitting the loss avoidance data. By constraining the plasticity parameters to one of 16 “quadrants,” consistent regions of parameter space redundancy were identified. (B) Illustrative examples of dopamine-weight change curves derived from quadrants which generated good fits (Q5 & Q15) to the control behaviour but biologically unrealistic changes (left panel D1 LTP occurred during a dopamine “dip”, right panel D2-LTD occurred during a dopamine “dip”). Solid lines represent the D2 curve, dashed lines the D1 curve. Quadrant 13, where the parameter space was bound by signs [+1 +1 −1−1] generated the most reliable fits with biologically feasible weight change curves (middle panel example).

The final plasticity parameter estimates for the controls (Figure 8) accurately reproduced controls subject’s behavior (Figure 3A), and dopamine–synaptic weight change curves were generated (Figure 3B), comparable to those observed by Gurney et. al., 2015, (their Figure 13).

**Figure 8:**
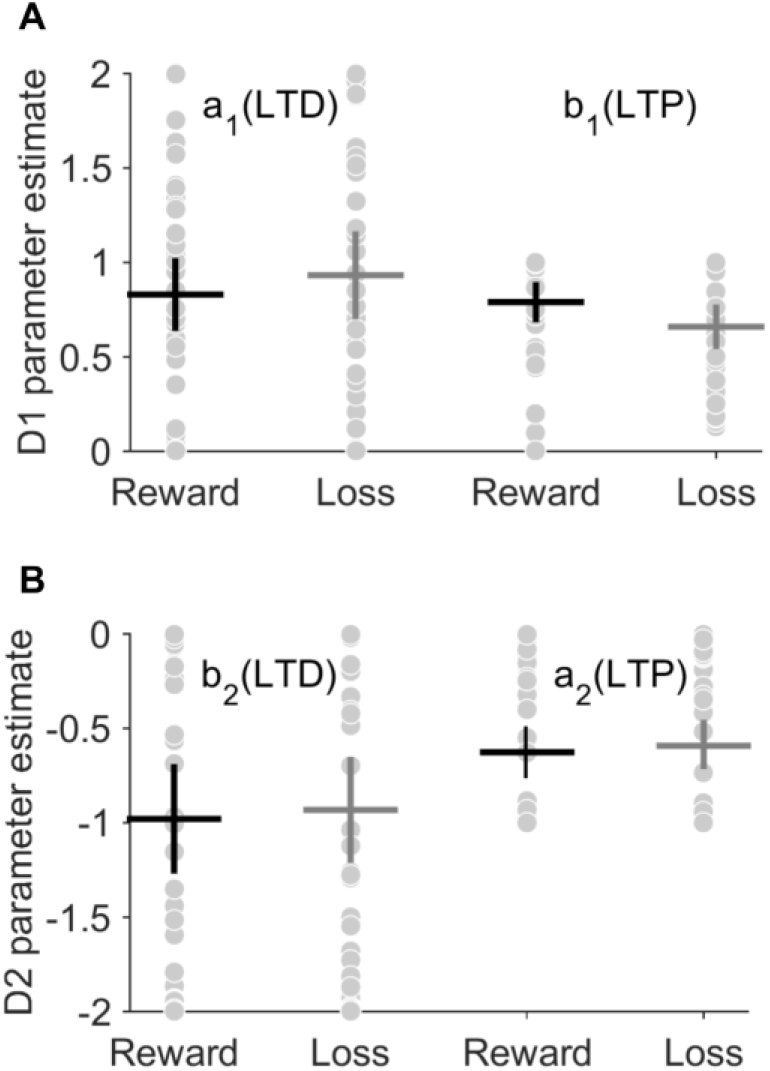
Basal ganglia model fitting recovers realistic plasticity parameter estimates for controls. Final plasticity parameter estimates for D1 in **(A)** and D2 **(B)** coefficients. Reward and Loss avoidance were fitted separately with individuals estimates represented by grey dots. Mean and 95% confidence limits represented by black and grey crosses for reward and loss respectively. The average plasticity parameter values were a_1_ = 0.83+/− 0.55, b_1_ = 0.78 +/− 0.27, a_2_= −0.97 +/− 0.85, b_2_ = −0.62 +/− 0.37 for reward and a_1_ = 0.93+/−0.67,b_1_ 0.65+/−0.31, a_2_ = −0.93+/− 0.82, b_2_ = −0.59 +/− 0.38 for the loss avoidance task.

#### 3.4.2 Patients

Previous *in vitro* studies of genetic forms of dystonia have demonstrated bidirectional abnormalities in cortico-striatal plasticity, including increases and decreases in LTP and LTD. To date it remains unknown if these are receptor specific or affect both D1 and D2 receptors. In order to test whether abnormal D1 (direct) or D2 (indirect) pathway cortico-striatal plasticity could explain the reward learning impairment in patients, four candidate hypothetical abnormalities of synaptic plasticity were tested: H_1_) increased D1-LTP : D1-LTD; H_2_) decreased D1-LTP : D1-LTP; H_3_) increased D2-LTP : D2-LTD; H_4_) decreased D2-LTP : D2-LTD. Each of these hypotheses were tested in turn by fitting the basal ganglia model to the patient’s behavioural data.

Comparing the negative log likelihood estimate for each hypothesis, H4 (decreased D2-LTP : D2-LTD) fitted the patient’s behavior best in 29 out of 40 patients for the reward task, with the next best hypothesis H1 (increased D1-LTP : D1-LTD) in 11 of 40, Chi-Square (1) = 14.45, p<0.001. Notably in the loss-avoidance task, despite no significant difference in behavioural performance, H4 (decreased D2-LTP: D2-LTD) also best explained patient’s behavior in 34 of 40 subjects, with the remaining 6 subjects best explained by H1 (increased D1-LTP : D1-LTD), Chi-Square (1) = 36.45, p<0.001. Thus, a single deficit of decreased D2-LTP: D2-LTD is consistent with both the patients’ worse reversal performance in the reward task and their reversal performance in the loss avoidance task.

Overlaying observed experimental choice behavior with the final best model fit demonstrated a close correspondence to both the reward (Figure 3A) and loss avoidance (Figure 9A), in both controls and patients. In contrast, there was a poor overlap between the modelled patients choice in both tasks when alternative plasticity hypotheses were fitted (H1-H3) - see Figure 10 (A & C).

**Figure 9:**
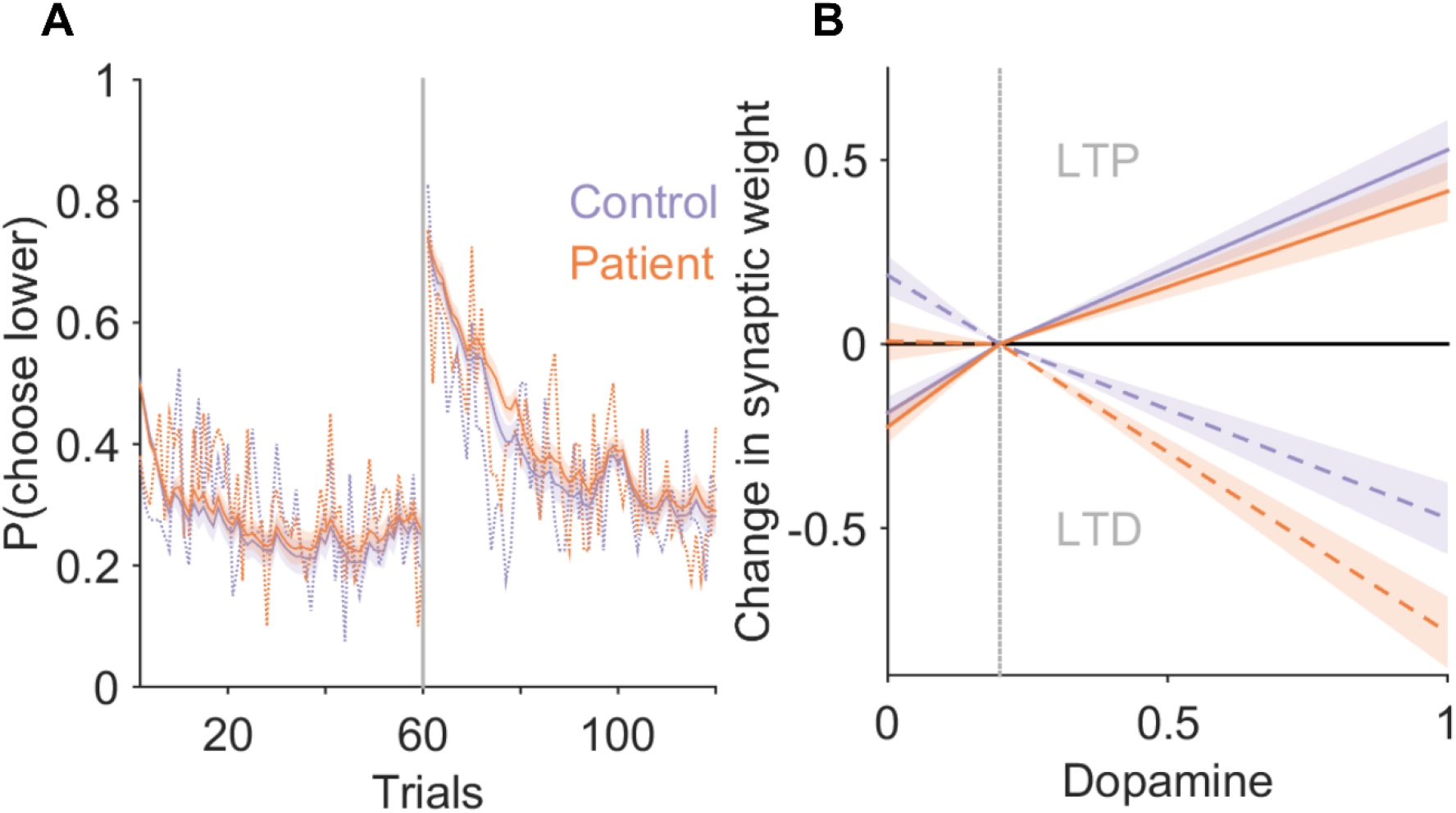
Loss avoidance learning in cervical dystonia is best explained by a model of the basal ganglia with abnormal D2 cortico-striatal plasticity. **(A)** Average probability of choosing low value actions “A1” and “A2” pre- and post-reversal reversal of contingencies in the reward task. Patient average choice probability in red, controls in blue. Vertical dotted line represents the point of contingency reversal. Dashed lines represent experimental behavior from patients and controls. The basal ganglia models performance following estimation of the optimal model parameters is superimposed (solid lines with 95% confidence limits represented by the shaded region). The models behavior closely overlaps with the experimental behavior of both the patients and controls. Despite no behavioural difference in the loss avoidance learning, a basal ganglia model with increased D2-LTD : LTP explained the patients loss avoidance strategy best in 31 of the 40 patients fitted. **(B)** The average dopamine-weight change curve for the patients (red) and controls (blue) with 95% confidence limits represented by the shaded region. Solid lines, represent D1 and dashed lines, represent D2 curves respectively.

**Figure 10:**
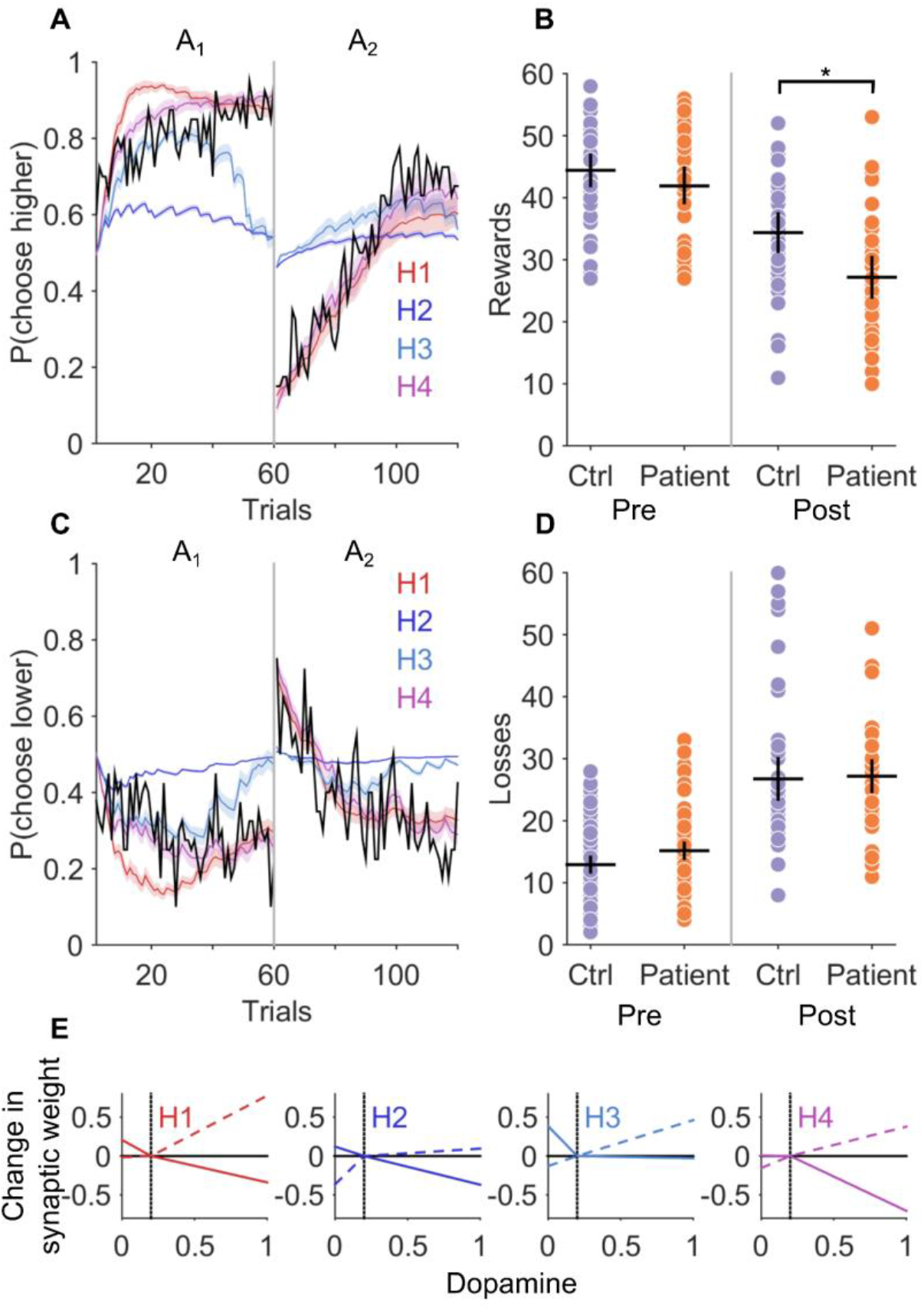
Model hypotheses comparison and generated behavior in reward and loss avoidance tasks. **(A)** and **(C)** the mean choice probability for the high and low value fractal for the patients is plotted in black. Superimposed are the result of fitting the basal ganglia model to each of the four hypothetical plasticity abnormalities (H1, D1 increased LTP : LTD; H2, D1 increased LTD : LTP; H3, increased D2 LTP : LTD; H4, increased D2 LTD : LTP). Average model choice probabilities with 95% confidence limits represented by the shaded area. **(B)** Behavioural data generated by the basal ganglia model performing the task with the “winning” hypothesis H4 (increased D2 LTD : LTP) reproduces the post reversal reward impairment (*p<0.01) with similar loss avoidance performance in patients and controls **(D). (E)** Final dopamine – weight change curves for each of the four hypothetical abnormalities in D1 or D2 cortico-striatal plasticity. Dashed lines D1, solid lines D2.

Using the final optimal parameter estimates for patients (using hypothesis H4) and controls, simulated behavioural data was generated using the basal ganglia model performing the original task. This simulated behaviour reproduced both the post-reversal impairment in the patients for the reward task (Figure 10B) and performance in the loss avoidance task (Figure 10D).

Overall, this analysis and results provides strong evidence in support of a pathological abnormality of D2 cortico-striatal plasticity, with excess synaptic depotentiation leading to a measureable impairment in reward reversal learning.

## 4.0 Discussion

This is the first study to demonstrate that patients with cervical dystonia have impaired reinforcement learning. A novel reinforcement learning basal ganglia model was developed which, when fitted to control subjects’ behaviour, successfully recovered measurable and biologically realistic synaptic parameters. Our results suggest that abnormal reversal learning in patients with cervical dystonia is best explained by abnormally high levels of D2 synaptic depotentiation. Notably, striatal D2 receptors are thought to be critical to reversal learning, as they are sensitive to the pause in dopamine release that accompanies trials following the reversal where reward is expected but not forthcoming (Frank, 2005; Cools *et al*., 2009; Cox *et al*., 2015). Selective blockade of D2 receptors in humans, or destruction of D2 expressing MSN’s in animals, leads to impairments in the reversal learning phase of reward learning tasks (Mehta *et al*., 2001; Nakanishi *et al*., 2014). Our reversal learning impairment in this group of patients is therefore consistent with these observations. Abnormal D2-receptor function is also consistent with SPECT studies reporting a selective reduction in striatal D2 receptor expression (Naumann *et al*., 1998) and the recognized clinical relationship between chronic D2 receptor blockade and tardive dystonia (Sethi, 2007).

Our modelling supports the idea that the reversal learning impairment in these patients could be caused by a relative absence of D2-LTP associated with the “dip” in dopamine particularly at the time of contingency reversal. The insufficient “NoGo” activity that would be a consequence of this and would result in a delay to suppress the decision to choose the opposite action with high pre-reversal value. Furthermore, our results predict that such NoGo activity required to suppress a previously learnt choice has to rise from an initially depotentiated state. This is a direct consequence of excessive D2-LTD that accompanies the “bursts” of dopamine associated with the pre-reversal acquisition phase of the task. This mechanism therefore suggests that perseveration errors that led to poorer task performance were mediated by impaired response inhibition, rather than over learning (increased LTP) or an inability to extinguish the previously learnt action values (impaired LTD). This distinction is important, as all of the four hypothesis tested could have provided equally plausible mechanistic explanations for the observed abnormal behavior.

Our results extend previous work (Arkadir *et al*., 2016) which demonstrated abnormal reward learning in patients with DYT1 generalized dystonia. In addition, our results support a more general conclusion, that abnormal reward based learning may be a common phenotypical abnormality in both generalized and common focal forms of dystonia. This parallels the original clinical observations of Marsden and Harrison(Marsden & Harrison, 1974), who realized the overlapping features between “spasmodic torticollis” and “idiopathic Torsion dystonia”.

There are some possible study limitations. Computational modeling cannot be used to make definitive statements about a synaptic level phenomena which is only directly accessible to *in vitro* neurophysiological techniques. Its crucial role is to generate hypotheses about disease mechanisms, which can be tested by experimental replication and manipulation of behavior based on the model’s predictions, and by interpretation of clinical trials of novel medications with known effects on synaptic plasticity. In the absence of a clearly defined genetic cause and corresponding animal model for the commonest forms of focal dystonia, it is unlikely that any other approach can be used to address the hypotheses we had for this study due to ethical considerations. As described previously in patients with cervical dystonia, we found higher levels of anxiety and depressive thoughts. These were at levels which were did not reach the threshold for being clinically significant for mild depression or anxiety and are unlikely to be responsible for the behavioral impairment observed as impaired reversal learning has only been observed in patients with severe depression or bipolar disorder (Dickstein *et al*., 2010).

In contrast to their established role in the treatment of generalized dystonia, anti-cholinergic medications have a highly variable and frequently limited role in the treatment of cervical dystonia (Nutt *et al*., 1984; Brans *et al*., 1996). The results reported here may go some way to explain this dichotomy as reduced cholinergic interneuron (TAN) firing is thought to lead to increased D2 synaptic depotentiation and impaired reward reversal learning (Franklin & Frank, 2015). In contrast, in animal models of generalized DYT1 dystonia, reduced cortico-striatal LTD is restored by anti-cholinergic’s (Sciamanna *et al*., 2012; Maltese *et al*., 2014). Our results suggest that if cholinergic abnormalities underpin abnormal cortico-striatal plasticity in cervical dystonia, then striatal cholinergic *hypofunction* is more likely in focal forms of the disease and would explain why cholinergic antagonism provides little symptom benefit. We therefore predict that drugs which reduce dopamine’s D2 inhibitory influence (eg. tetrabenazine or valbenazine) on cholinergic interneuron excitability, or increase cholinergic interneuron excitability directly (M1 receptor agonists), would improve reversal learning in cervical dystonia, and may ultimately have a therapeutic role in the treatment of dystonia. For the third of patients who receive no benefit from existing treatments (Misra *et al*., 2012), future studies which combine reward learning tasks with a single dose pharmacological challenge may be used to identify new agents, based on their ability to improve reward learning behavior. Furthermore, this approach could be used to stratify subgroups of patients according to their response, so that personalized treatments with disease modifying potential can be selected for future clinical trials.

## 5.0 Conclusions

In summary, we report a behavioral reversal learning deficit in patients with cervical dystonia, extending previous work on DYT1 dystonia. Abnormal reversal learning behaviour was best explained by decreased D2-LTP, suggesting excessive D2 cortico-striatal synaptic depotentiation. Computational modelling of behavior in patients can be used to test hypotheses of abnormal synaptic plasticity inaccessible by other means and thus to interpret the behavioral effects of novel treatments.

## Acknowledgements

Funded by a Dystonia Society (UK) grant to TG and MDH funded by a MRC Senior non-Clinical Fellowship (MR/J008648/1). Dr Blair Johnston wrote an earlier version of the code which presents the cognitive task. The authors would like to thank Dr Kerr Grieve and Professors Peter Brown and Miratul Muqit for comments on earlier version of the manuscript. In addition, we would like to thank the patients who participated and Dr’s Carl Counsell, Louise Davidson and Edward Newman for their assistance with patient recruitment.

## Author Contributions

TG and DS designed research. TG and MH contributed to analytical tools and performed analysis of the data. TG, DS and MH wrote the paper.

## Data accessibility

The code and behavioural data can be requested by email to the corresponding author (TG).

## Abbreviations

LTP: Long term synaptic potentiation
LTD: Long term synaptic depression
RPE: Reward prediction error
MNI: Montreal Neurological Institute
CTRL: Control
CD: Cervical dystonia

## Supporting Information

**Supplementary Figure 1:-.**
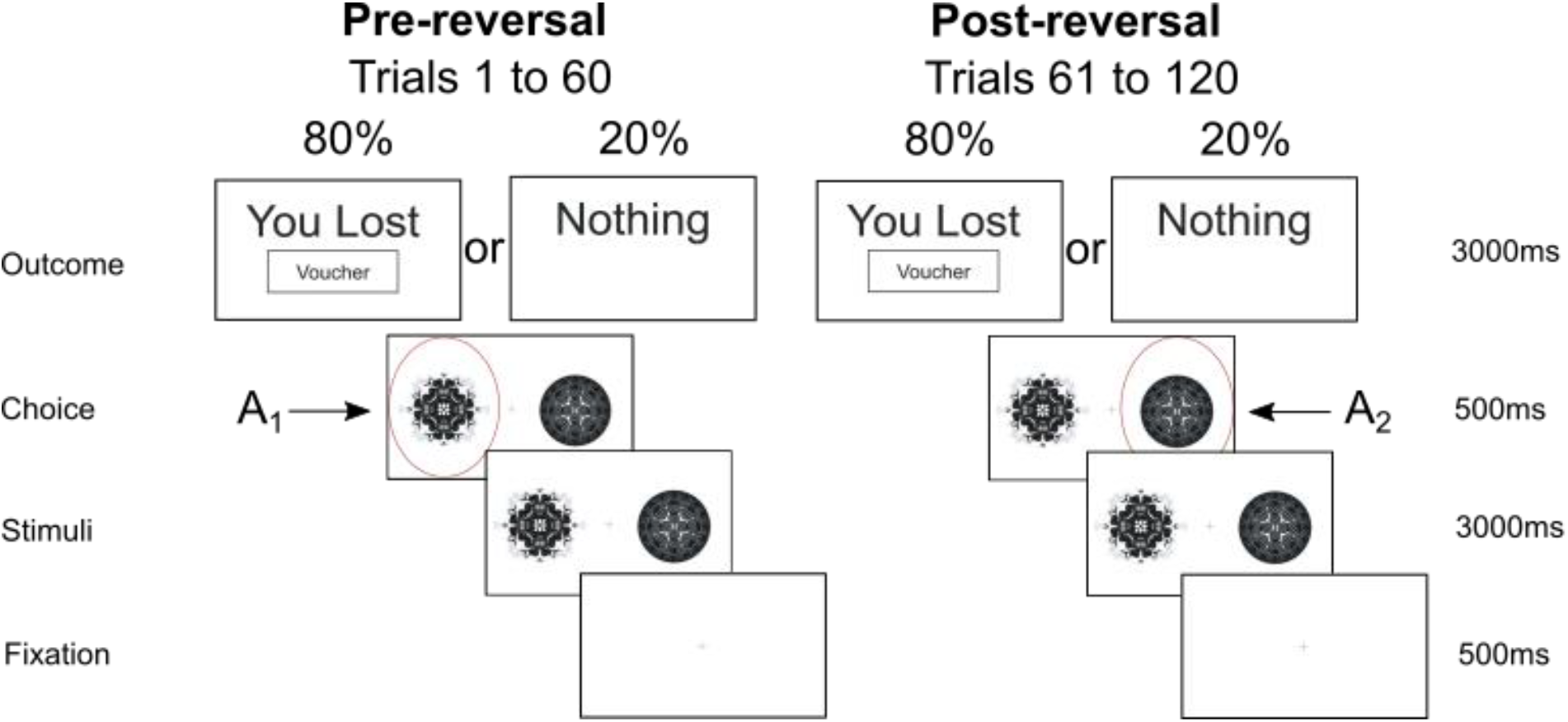
Probabilistic reversal learning task. Example of fractal images presented during a single loss trial. The probability of receiving a loss “voucher” reverses after 60 trials requiring the participant to suppress the previously learnt choice and learn to “reverse” their decision to choose the previously low value (pre-reversal) fractal. In the pre-reversal phase choice of A_1_ is associated with an 80% outcome of loss. Post-reversal this reduces to 20% with the choice of A_2_ being associated with 80%.

